# A key spectral tuning site of UV-sensitive vertebrate non-visual opsin Opn5

**DOI:** 10.1101/2025.05.19.654871

**Authors:** Takahiro Yamashita, Kazuyuki Asamoto, Kengo Fujii, Chihiro Fujiyabu, Hideyo Ohuchi, Yoshinori Shichida

**Author notes:** Corresponding author: Takahiro Yamashita, Department of Biophysics, Graduate School of Science, Kyoto University, Kyoto 606-8502, Japan.

## Abstract

Opsins are photoreceptive proteins responsible for visual and non-visual photoreceptions in animals. In general, vertebrates have multiple visual and non-visual opsins whose spectral sensitivities range from the UV to the red region. Among these opsins, Opn5 has been widely identified in vertebrates from fishes to primates and functions as a non-visual opsin in various tissues including the retina and brain. Vertebrate Opn5 has been characterized as a UV-sensitive bistable opsin. Thus, Opn5 provides one of the molecular mechanisms determining the short wavelength limit that vertebrates can detect. In this study, we searched for the amino acid residue responsible for the UV light sensitivity of Opn5. Our mutational analysis revealed that Opn5 acquired visible light sensitivity by the substitution of Lys91 with an amino acid other than arginine or tyrosine residue. In addition, the mutations at Lys91 altered the preferential binding of the retinal isomers in Opn5. Therefore, the conservation of Lys91 among vertebrate Opn5 proteins would be necessary to enable Opn5 to work as the shortest wavelength sensor in various tissues.

## Introduction

Animals utilize light as an important information source for various physiological functions. Opsins are universal photoreceptive proteins for both visual and non-visual photoreceptions in animals and are characterized as photosensitive G protein-coupled receptors. Recent accumulation of genomic information revealed that animals have various kinds of opsin genes, which are classified into several groups based on their sequences (1, 2). All opsins share common structural elements including seven transmembrane domains and bind a light-absorbing chromophore, retinal, via a Schiff base linkage to Lys296 (based on the bovine rhodopsin numbering system) of the protein moiety. Among these opsins, vertebrate rhodopsin is the best-studied opsin (3). Vertebrate rhodopsin functions in rod cells of the retina for visual photoreception and binds 11-*cis* retinal in the dark. After photoreception, the isomerization of the retinal to all-*trans* form induces the formation of the meta-stable active state, which triggers G protein signaling and eventually evokes the hyperpolarization of rod cells.

In the human genome, we can find nine opsin genes, that is, four visual opsins (rhodopsin and three cone visual pigments), Opn3, Opn4, Opn5, Rgr and Rrh (peropsin) (4). Among these opsins, Opn5 is the most recently identified opsin and forms an independent opsin group in the opsin classification (5). Opn5 genes have been identified in a wide range of vertebrate species from fishes to primates and these vertebrate Opn5 genes are classified into several subgroups (4, 6, 7). Non-mammalian vertebrates have multiple Opn5 genes, whereas most mammals, including human and mouse, have only one Opn5 gene (Opn5m). Previous studies revealed that Opn5m proteins of various vertebrate species from fishes to human bind 11-*cis* retinal to form UV light-sensitive photo-pigments (8, 9). UV light irradiation of these pigments induces isomerization of the retinal to the all-*trans* form to produce a visible light-sensitive active state. This active state is thermally stable and photo-converts back to the original dark state. Thus, Opn5m has two stable states, the dark and active states, which are inter-convertible with each other by photoreception. This means that Opn5m is categorized as a bistable opsin. In addition, our analysis of the distribution patterns detected the expression signals of Opn5m in a subset of the ganglion cells of the retina and the preoptic area of the hypothalamus in mouse and common marmoset (9). (9). Recent studies using Opn5m knock-out mice have advanced the understanding of the physiological relevance of Opn5m. Absorption of short wavelength light by Opn5m contributes to the entrainment of the circadian rhythm in several tissues (10–13), the regulation of vascular development and choroidal thickness in the eyes (14, 15) and the suppression of thermogenesis (16). (16). In non-mammalian vertebrates, it has also been revealed that Opn5m works for several important non-visual photoreceptive functions. In avian species, Opn5m is expressed in the paraventricular organ of the hypothalamus and controls the photoperiodic induction of testicular growth (8, 17). In medaka fish, Opn5m is expressed in the pituitary gland and regulates the secretion of melanocyte-stimulating hormone to change the black pigmentation in the body (18, 19). Thus, Opn5m underlies a variety of the short wavelength light-dependent biological responses in vertebrates.

Opsins have their characteristic spectral sensitivities ranging from the UV to the red region. Among the human opsins, Opn5m is considered to be the opsin that is sensitive to the shortest wavelength light (9). In addition, the conservation of UV light sensitivity of Opn5m across vertebrate species suggests that Opn5m provides one of the molecular bases underlying the absorption of the shortest wavelength light in vertebrates (8, 9). In this context, we searched for the amino acid residue(s) responsible for the UV light sensitivity of vertebrate Opn5m. In general, visible light-sensitive opsins bind the retinal via a protonated Schiff base linkage, whereas UV light-sensitive opsins bind it via a deprotonated Schiff base linkage (1, 3). Thus, the spectral difference between visible light-sensitive and UV light-sensitive opsins is mainly regulated by the protonation state of the Schiff base. Previous studies of UV light-sensitive opsins revealed that the amino acid residues in the extracellular side of Helix II, which are speculated to be located in the vicinity of the Schiff base in the 3D structures of the opsins, are responsible for the UV light sensitivity of the opsins (20–25). Here, our mutational analysis showed that Lys91 in the extracellular side of Helix II is well conserved in vertebrate Opn5m and is necessary for the UV light sensitivity of Opn5m. Moreover, we also found that this lysine residue controls the ability of Opn5m to directly bind the retinal isomers. Based on comprehensive mutational analysis at this site, we would like to discuss the molecular mechanism by which Opn5m works as the shortest wavelength sensor in various tissues such as retina and brain of vertebrates.

## Results

### Search for the amino acid residue(s) responsible for UV light sensitivity

Our previous studies revealed that not only mammalian Opn5 but also non-mammalian Opn5m forms a UV-sensitive bistable opsin (8, 9). Moreover, in theose studies, *Xenopus tropicalis* Opn5m showed the highest expression yield in cultured cells among various Opn5m recombinant proteins that we analyzed. Wild-type *X. tropicalis* Opn5m directly binds all-*trans* retinal to produce a visible light-absorbing form (Fig. 1A). Yellow light irradiation induces the formation of a UV light-absorbing form, and subsequent UV and yellow light irradiations result in inter-conversion between the UV light- and visible light-absorbing forms. These spectral changes are triggered by the photoisomerization of the retinal between 11-*cis* and all-*trans* forms (8), which enables calculation of the absorption spectra of the 11-*cis* retinal and all-*trans* retinal bound forms of Opn5m (Fig. 1B). In this study, we performed mutational analysis of *X. tropicalis* Opn5m to search for the amino acid residue(s) responsible for the UV light sensitivity. The spectral tuning of UV-sensitive opsins has been extensively analyzed in vertebrate cone visual pigments and insect visual opsins. Previous studies revealed the importance of the amino acid residues in the extracellular side of Helix II for the spectral tuning of UV-sensitive opsins (20–25). S90C mutation of chicken violet-sensitive cone pigment resulted in UV light sensitivity of the pigment, whereas C90S mutation of zebra finch UV-sensitive cone pigment resulted in violet light sensitivity of the pigment (25). In addition, F86Y mutation of mouse UV-sensitive cone pigment produced a violet light-sensitive pigment, whereas Y86F mutation of bovine violet-sensitive cone pigment produced a UV light-sensitive pigment (20). These results suggested that the introduction of amino acids whose side chain contained a hydroxyl group at positions 86 and 90 led to the acquisition of visible light sensitivity. The analysis of Drosophila opsins showed that K90 mutants of UV-sensitive opsins formed violet-sensitive pigments, which suggested the importance of the lysine residue for the UV light sensitivity (22, 23). The comparison of the sequences in the extracellular side of Helix II among vertebrate Opn5m highlighted the conservation of Lys91 in this region (Fig. S1A). Based on these previous studies, we prepared five mutants (I86S, I86T, G90S, G90T and K91T) of Opn5m (Fig. S1B) and analyzed their spectral property.

**Figure 1.**
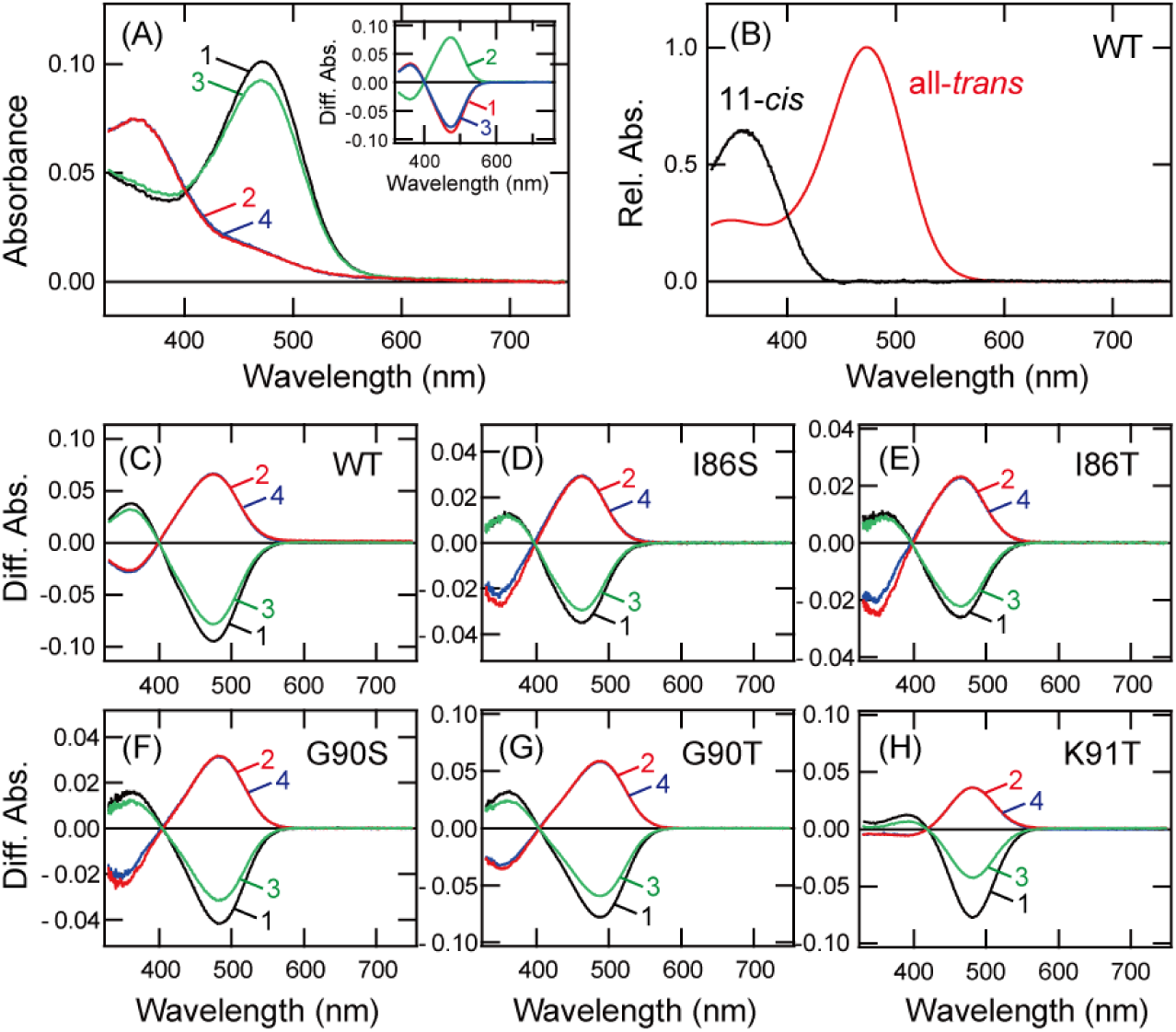
Spectral property of wild-type and mutants of Xenopus Opn5m. (A) Absorption spectra of wild-type Opn5m purified after incubation with all-*trans* retinal. Spectra were recorded in the dark (curve 1), after yellow light (>500 nm) irradiation (curve 2), after subsequent UV light (360 nm) irradiation (curve 3) and after yellow light re-irradiation (curve 4). (inset) Spectral changes of wild-type Opn5m caused by yellow light irradiation (curve 1), subsequent UV light irradiation (curve 2) and yellow light re-irradiation (curve 3). (B) Calculated absorption spectra of wild-type Opn5m. The method of calculating the spectra of 11-*cis* (black curve) and all-*trans* retinal (red curve) bound forms of Opn5m is described in previous papers (8, 9). The spectrum of 11-*cis* retinal or all-*trans* retinal bound forms has a peak in the UV region (360 nm) or in the blue region (474 nm), respectively. (C-H) Spectral changes of detergent-solubilized cell membranes expressing Opn5m. The cell membranes containing wild-type (C), I86S (D), I86T (E), G90S (F), G90T (G) and K91T (H) after the addition of all-*trans* retinal were solubilized with 1% DDM and their absorption spectra were recorded in the dark and after light irradiation. Spectral changes caused by yellow light irradiation (curve 1), subsequent UV light irradiation (curve 2), yellow light re-irradiation (curve 3) and UV light re-irradiation (curve 4) are shown.

We obtained the cell membranes containing the wild-type and five mutants of Opn5m after the addition of all-*trans* retinal and solubilized them with 1% dodecyl maltoside (DDM). Yellow light irradiation of the wild-type and four of the mutants (I86S, I86T, G90S and G90T) resulted in highly similar spectral changes, namely, a decrease of absorbance in the blue region and a concomitant increase of absorbance in the UV region (Fig. 1C-1G). Spectral changes induced by subsequent UV and yellow light irradiations were mirror images of each other in these samples. Detailed comparison of the spectral change showed that the negative maximum, which corresponds to the peak of the all-*trans* retinal bound form, was slightly red-shifted in G90 mutants and slightly blue-shifted in I86 mutants (Fig. S2). These results indicated that I86S, I86T, G90S and G90T mutants are UV-sensitive bistable opsins, like wild-type, and these mutations had a slight effect on the spectral peak of the all-*trans* retinal bound form. By contrast, yellow light irradiation of K91T mutant resulted in a quite small increase of absorbance in the UV region accompanied by a large decrease of absorbance in the visible region (Fig. 1H). The spectral changes induced by subsequent UV and yellow light irradiations in K91T mutant were mirror images of each other. This result suggested that K91T mutation substantially affected the spectral property of Opn5m without hindering the bistable photoreaction.

### Molecular property of K91T mutant

To analyze the detailed spectral property of K91T mutant, we purified K91T mutant after the addition of 11-*cis* retinal (Fig. 2A). The absorption spectrum had a peak at around 470 nm and UV light irradiation decreased the absorbance in the blue region. Subsequent yellow light irradiation induced a decrease of absorbance in the blue region and a quite small increase of absorbance in the UV region. The analysis of the retinal configuration revealed that the dark state predominantly contained all-*trans* retinal, which is formed by thermal isomerization of 11-*cis* retinal in culture medium, and yellow and UV light irradiations triggered the photoisomerization of the retinal into the 11-*cis* form (Fig. S3A). We also purified K91T mutant after the addition of all-*trans* retinal and analyzed the spectral and retinal configuration changes by light irradiations (Figs. 2B and S3B). In the dark, K91T mutant sample had a spectral peak at around 470 nm and predominantly contained all-*trans* retinal. Yellow light irradiation led to a decrease of absorbance in the blue region by the conversion from all-*trans* retinal to 11-*cis* retinal, and subsequent UV light irradiation resulted in some recovery of absorbance in the blue region by the conversion of the retinal into all-*trans* form. Yellow and UV re-irradiations induced repetitive spectral changes, which indicated that K91T mutant retained the bistable photoreaction. Based on the results shown in Figs. 2B and S3B, we calculated the absorption spectra of 11-*cis* and all-*trans* retinal bound forms of K91T mutant (Figs. 2C and S3C). K91T mutation induced an ∼75 nm red shift of the absorption maximum (λmax) in the 11-*cis* retinal bound form (435 nm) to form a visible light-sensitive pigment. By contrast, K91T mutation had only a slight effect on λmax of the all-*trans* retinal bound form (472 nm). Moreover, K91T mutant exhibited preferential binding to all-*trans* retinal.

**Figure 2.**
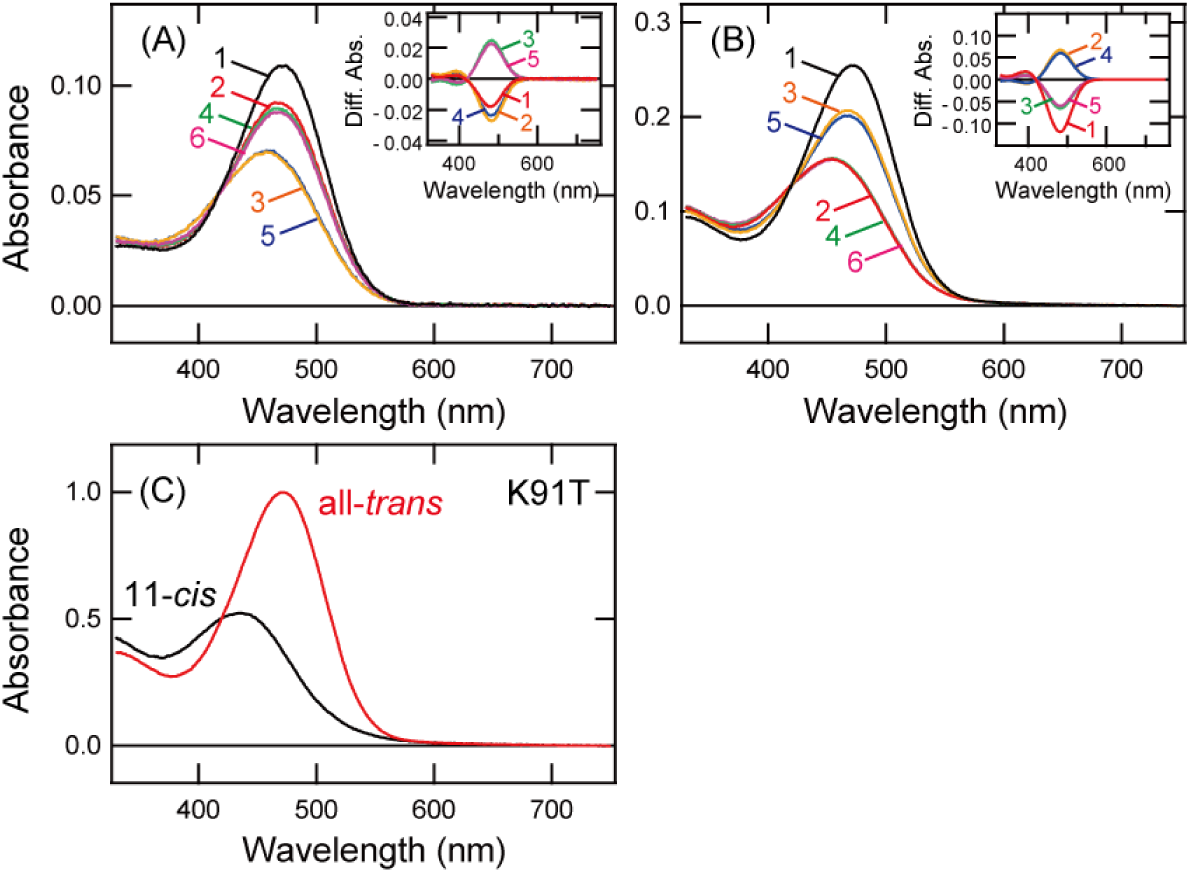
Spectral property of K91T mutant of Opn5m. (A) Absorption spectra of K91T mutant of Opn5m purified after the incubation with 11-*cis* retinal. Spectra were recorded in the dark (curve 1), after UV light (360 nm) irradiation (curve 2), after subsequent yellow light (>500 nm) irradiation (curve 3), after UV light re-irradiation (curve 4), after yellow light re-irradiation (curve 5) and after UV light re-irradiation (curve 6). (inset) Spectral changes of K91T mutant caused by UV light irradiation (curve 1), subsequent yellow light irradiation (curve 2), UV light re-irradiation (curve 3), yellow light re-irradiation (curve 4) and UV light re-irradiation (curve 5). (B) Absorption spectra of K91T mutant of Opn5m purified after the incubation with all-*trans* retinal. Spectra were recorded in the dark (curve 1), after yellow light irradiation (curve 2), after subsequent UV light irradiation (curve 3), after yellow light re-irradiation (curve 4), after UV light re-irradiation (curve 5) and after yellow light re-irradiation (curve 6). (inset) Spectral changes of K91T mutant caused by yellow light irradiation (curve 1), subsequent UV light irradiation (curve 2), yellow light re-irradiation (curve 3), UV light re-irradiation (curve 4) and yellow light re-irradiation (curve 5). (C) Calculated absorption spectra of K91T mutant of Opn5m. λmax of the 11-*cis* retinal and all-*trans* retinal bound forms are 435 nm and 472 nm, respectively. The method of calculating the spectra of 11-*cis* (black curve) and all-*trans* retinal (red curve) bound forms of K91T mutant is provided in Fig. S3.

### Spectral property of other K91 mutants

The mutational analysis on the extracellular side of Helix II suggested the possibility that the well-conserved lysine residue at position 91 is a spectral tuning site in UV-sensitive Opn5m. Next, we analyzed the effects of other mutations at Lys91 on the spectral property of Opn5m. We replaced Lys91 with the other 18 naturally occurring amino acid residues and prepared the cell membranes containing these mutants after the addition of all-*trans* retinal (Fig. S4). After solubilizing the cell membranes with 1% DDM, we observed substantial spectral changes caused by light irradiations. Thus, all of these mutants formed photo-pigments upon the reconstitution with retinal.

Comparison of the spectral changes showed that, in all the mutants except K91R mutant, yellow light irradiation induced a large decrease of absorbance in the visible region, as in the wild-type (Fig. 1C). This suggested that K91R mutant has lost the ability to directly bind all-*trans* retinal. In addition, K91Y mutant exhibited an increase of absorbance in the UV region after yellow light irradiation, as did the wild-type (Fig. 1C), whereas the other mutants exhibited quite a small increase of absorbance in the UV region after yellow light irradiation, as did K91T mutant (Fig. 1H). We also prepared the cell membranes containing these mutants after the addition of 11-*cis* retinal (Fig. S5). After the cell membranes were solubilized with 1% DDM, K91R exhibited a decrease of absorbance in the UV region and a concomitant increase of absorbance in the blue region after UV light irradiation. This suggests that K91R mutant maintains the ability to directly bind 11-*cis* retinal. Moreover, in many of these mutants (K91A, K91C, K91D K91F, K91G, K91I, K91N, K91P, K91S, K91V and K91W), as in K91T mutant (inset of Fig. 2A), UV light irradiation did not induce a substantial increase of absorbance in the blue region. This suggested that these mutants have decreased affinity for 11-*cis* retinal. Based on these spectral analyses, we speculated that K91R and K91Y mutants form UV light-sensitive pigments like wild-type, whereas the other mutants form visible light-sensitive pigments like K91T mutant.

### Molecular property of K91A and K91Q mutants

Next, we analyzed the detailed spectral property of several mutants. Among the 18 mutants shown in Figs. S4 and S5, we purified K91A and K91Q mutants, which are considered to form visible light-sensitive pigments, after the addition of 11-*cis* retinal (Figs. 3A and 3C). These mutant samples had spectral peaks in the blue region and contained not only 11-*cis* retinal but also a substantial amount of all-*trans* retinal (Figs. S6A and S6C). In K91A mutant, UV light irradiation and subsequent yellow light irradiation caused the isomerization of all-*trans* retinal to 11-*cis* retinal and decreased the absorbance in the blue region (Figs. 3A and S6A). In K91Q mutant, UV light irradiation increased the absorbance in the blue region by the isomerization of 11-*cis* retinal to all-*trans* retinal and subsequent yellow light irradiation decreased the absorbance in the blue region by the isomerization of all-*trans* retinal to 11-*cis* retinal (Figs. 3C and S6C). Moreover, we prepared purified samples of K91A and K91Q mutants after the addition of all-*trans* retinal (Figs. 3B and 3D). These mutant samples had spectral peaks in the blue region and predominantly contained all-*trans* retinal (Figs. S6B and S6D). Yellow light irradiation of these samples decreased the absorbance in the blue region and subsequent UV light irradiation resulted in some recovery of absorbance in the blue region. These spectral changes occurred as a result of photo-conversion between 11-*cis* retinal and all-*trans* retinal. Based on these results, we calculated the absorption spectra of the 11-*cis* retinal and all-*trans* retinal bound forms of K91A and K91Q mutants (Figs. 3E and 3F). As speculated based on the results shown in Figs. S4 and S5, the 11-*cis* retinal bound forms of K91A and K91Q mutants were maximally sensitive to violet light (λmax at 428 nm and 424 nm, respectively). By contrast, K91A and K91Q mutations had a quite small effect on the spectral sensitivity of the all-*trans* retinal bound form. Moreover, we confirmed that K91A and K91Q mutations resulted in preferential binding to all-*trans* retinal. The spectral property and binding preference of retinal isomers of these mutants are similar to those of K91T mutant (Fig. 2).

**Figure 3.**
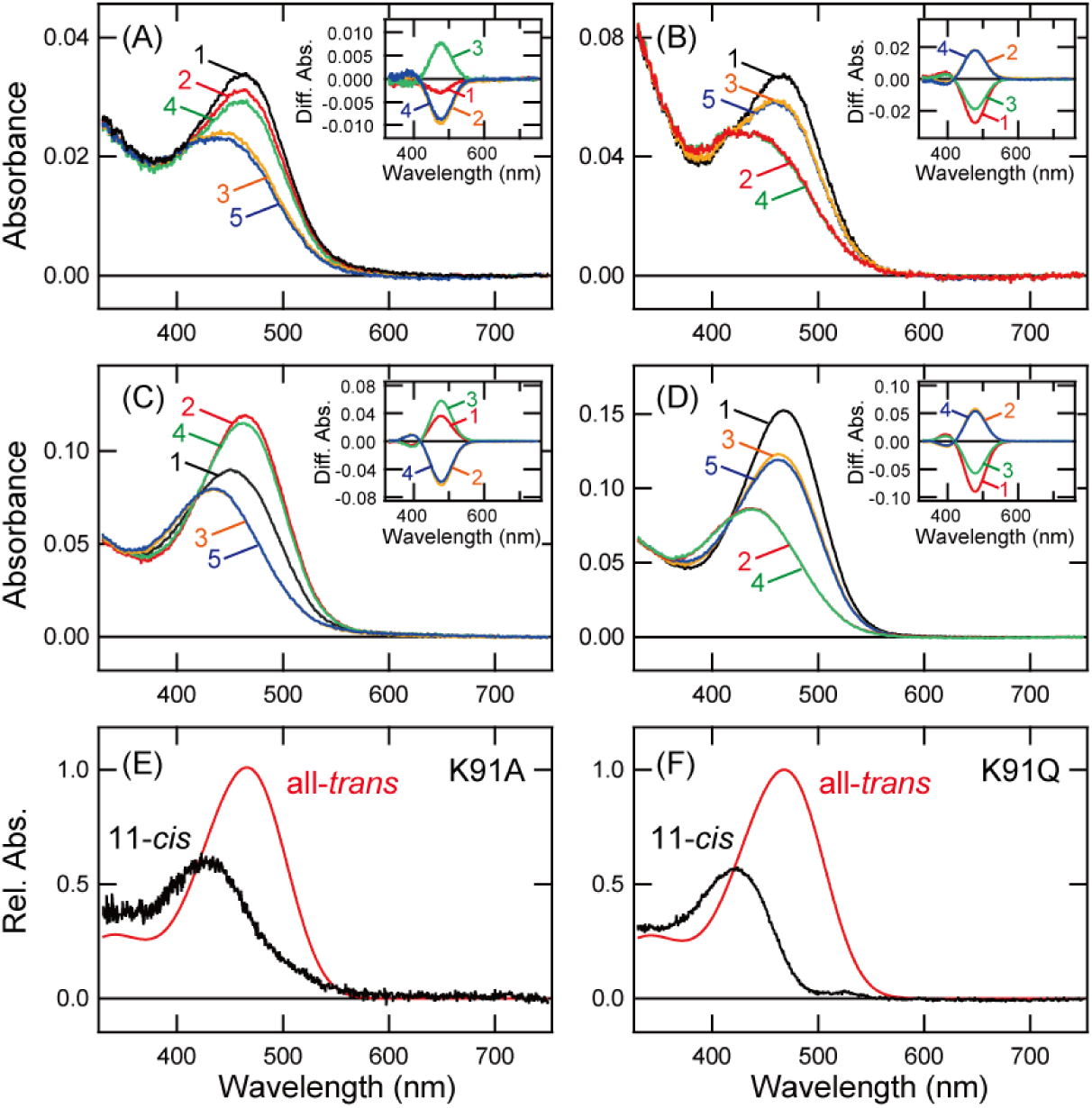
Spectral property of K91A and K91Q mutants of Opn5m. (A, C) Absorption spectra of K91A (A) and K91Q (C) mutants of Opn5m purified after the incubation with 11-*cis* retinal. Spectra were recorded in the dark (curve 1), after UV light (360 nm) irradiation (curve 2), after subsequent yellow light (>500 nm) irradiation (curve 3), after UV light re-irradiation (curve 4) and after yellow light re-irradiation (curve 5). (inset) Spectral changes caused by UV light irradiation (curve 1), subsequent yellow light irradiation (curve 2), UV light re-irradiation (curve 3) and yellow light re-irradiation (curve 4). (B, D) Absorption spectra of K91A (B) and K91Q (D) mutants of Opn5m purified after the incubation with all-*trans* retinal. Spectra were recorded in the dark (curve 1), after yellow light irradiation (curve 2), after subsequent UV light irradiation (curve 3), after yellow light re-irradiation (curve 4) and after UV light re-irradiation (curve 5). (inset) Spectral changes caused by yellow light irradiation (curve 1), subsequent UV light irradiation (curve 2), yellow light re-irradiation (curve 3) and UV light re-irradiation (curve 4). (E, F) Calculated absorption spectra of K91A (E) and K91Q (F) mutants of Opn5m. λmax of the 11-*cis* retinal and all-*trans* retinal bound forms of K91A are 428 nm and 466 nm, respectively. λmax of the 11-*cis* retinal and all-*trans* retinal bound forms of K91Q are 424 nm and 468 nm, respectively. The method of calculating the spectra of 11-*cis* (black curve) and all-*trans* retinal (red curve) bound forms is provided in Fig. S6.

### Molecular property of K91R and K91Y mutants

We also analyzed the molecular property of K91R and K91Y mutants, because these mutants are considered to form UV light-sensitive pigments. We purified these mutants after the addition of 11-*cis* retinal (Figs, 4A and 4C). K91R mutant sample had a spectral peak in the UV region (Fig. 4A) and predominantly contained 11-*cis* retinal (Fig. S7A). UV light irradiation shifted the spectrum into the blue region and subsequent yellow light irradiation recovered the spectrum in the UV region (Fig. 4A). These spectral changes were triggered by the photo-conversion between 11-*cis* retinal and all-*trans* retinal (Fig. S7A). K91Y mutant sample had a substantial absorbance in the blue region (Fig. 4C). This was probably because this sample contained a substantial amount of all-*trans* retinal in addition to 11-*cis* retinal (Fig. S7B). UV light irradiation and subsequent yellow light irradiation induced an increase and a decrease of absorbance in the blue region, respectively (Fig. 4C).

**Figure 4.**
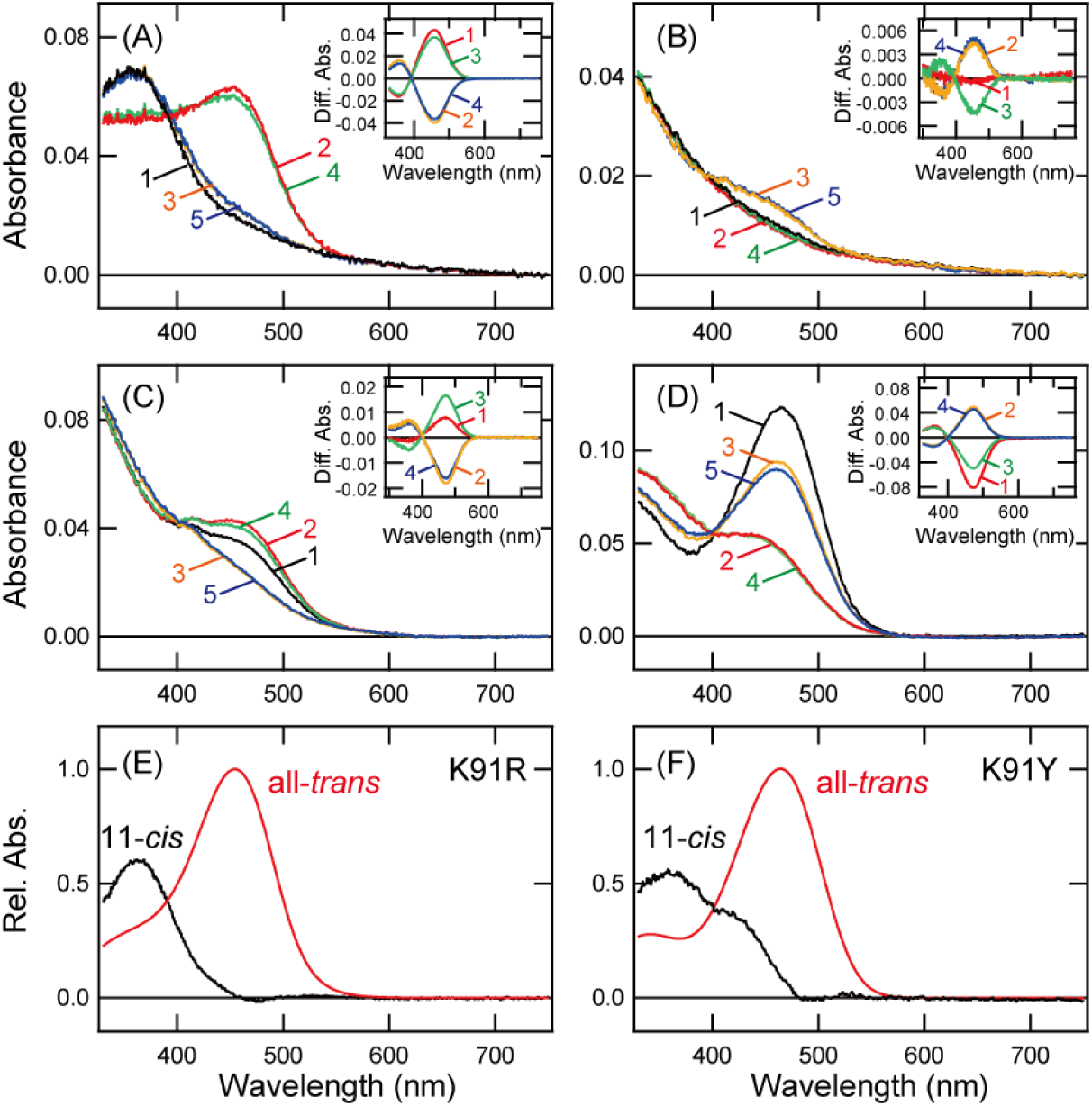
Spectral property of K91R and K91Y mutants of Opn5m. (A, C) Absorption spectra of K91R (A) and K91Y (C) mutants of Opn5m purified after the incubation with 11-*cis* retinal. Spectra were recorded in the dark (curve 1), after UV light (360 nm) irradiation (curve 2), after subsequent yellow light (>500 nm) irradiation (curve 3), after UV light re-irradiation (curve 4) and after yellow light re-irradiation (curve 5). (inset) Spectral changes caused by UV light irradiation (curve 1), subsequent yellow light irradiation (curve 2), UV light re-irradiation (curve 3) and yellow light re-irradiation (curve 4). (B, D) Absorption spectra of K91R (B) and K91Y (D) mutants of Opn5m purified after the incubation with all-*trans* retinal. Spectra were recorded in the dark (curve 1), after yellow light irradiation (curve 2), after subsequent UV light irradiation (curve 3), after yellow light re-irradiation (curve 4) and after UV light re-irradiation (curve 5). (inset) Spectral changes caused by yellow light irradiation (curve 1), subsequent UV light irradiation (curve 2), yellow light re-irradiation (curve 3) and UV light re-irradiation (curve 4). (E, F) Calculated absorption spectra of K91R (E) and K91Y (F) mutants of Opn5m. λmax of the 11-*cis* retinal and all-*trans* retinal bound forms of K91R are 362 nm and 454 nm, respectively. λmax of the 11-*cis* retinal and all-*trans* retinal bound forms of K91Y are 360 nm and 465 nm, respectively. The method of calculating the spectra of 11-cis (black curve) and all-trans retinal (red curve) bound forms is provided in Fig. S7.

These spectral changes were mainly caused by the inter-conversion between 11-*cis* retinal and all-*trans* retinal (Fig. S7B). We also purified these mutants after the addition of all-*trans* retinal (Figs. 4B and 4D). K91R mutant sample exhibited no clear spectral peak. Yellow light irradiation induced no spectral change and subsequent UV light irradiation induced only a quite small increase in the blue region (Fig. 4B), which is consistent with the result shown in Fig. S4. This spectral property indicated that K91R mutant lost the ability to directly bind all-*trans* retinal. K91Y mutant had a spectral peak in the blue region (Fig. 4D) and predominantly contained all-*trans* retinal (Fig. S7C). Yellow light irradiation shifted the spectrum into the UV region and subsequent UV light irradiation resulted in some recovery of absorbance in the blue region (Fig. 4D). During these spectral changes, inter-conversion between 11-*cis* retinal and all-*trans* retinal was observed (Fig. S7C). Based on these results, we calculated the absorption spectra of the 11-*cis* retinal and all-*trans* retinal bound forms of K91R and K91Y mutants (Figs. 4E and 4F). As speculated based on the results shown in Figs. S4 and S5, the 11-*cis* retinal bound form of K91R mutant was maximally sensitive to UV light (λmax at 362 nm) like wild-type (Fig. 4E). The all-*trans* retinal bound form of K91R mutant had a spectral peak at 454 nm, ∼20 nm blue-shifted from that of wild-type. Thus, K91R mutation resulted in maintenance of UV light sensitivity and preferential binding to 11-*cis* retinal. By contrast, the 11-*cis* retinal bound form of K91Y mutant had a spectral peak in the UV region and a spectral shoulder at around 430 nm (Fig. 4F). K91Y mutation had a small effect (∼9 nm blue-shift) on the spectral sensitivity of the all-*trans* retina bound form. This spectral property showed that K91Y mutant contained UV light-sensitive and visible light-sensitive components in the 11-*cis* retinal bound form.

Our spectral analysis was performed at pH 6.5. In this pH condition, a unique mixture of UV light-sensitive and visible light-sensitive components was observed in K91Y mutant. These two spectrally distinguishable components may result from the protonation and deprotonation of the Schiff base within the protein. Thus, we analyzed the spectral property of K91Y mutant under different pH conditions. K91Y mutant purified after the addition of all-*trans* retinal had a spectral peak in the blue region at pH 6 (Fig. 5A) and pH 7 (Fig. 5B). These two samples predominantly contained all-*trans* retinal (Fig. S8A and S8B). Yellow light irradiation of these two samples decreased the absorbance in the blue region and increased the absorbance in the UV region (Figs. 5A and 5B). In this process, the spectral change in the UV region at pH 7 was larger than that at pH 6 (insets of Figs. 5A and 5B). Moreover, the profiles of the retinal configuration changes caused by this yellow light irradiation were quite similar to each other (Figs. S8A and S8B). Based on these results, we calculated the absorption spectra of 11-*cis* retinal and all-*trans* retinal bound forms of K91Y mutant at pH 6 and pH 7 (Figs. 5C and 5D). Comparison of these spectra of the 11-*cis* retinal bound form showed that the ratio of the UV light-sensitive component at pH 7 was larger than that at pH 6. This can be explained by the increase of the ratio of the component with the deprotonated Schiff base at pH 7 compared to that at pH 6. Thus, these results suggested that K91Y mutant contained two spectrally distinguishable components which have protonated and deprotonated Schiff base under neutral conditions.

**Figure 5.**
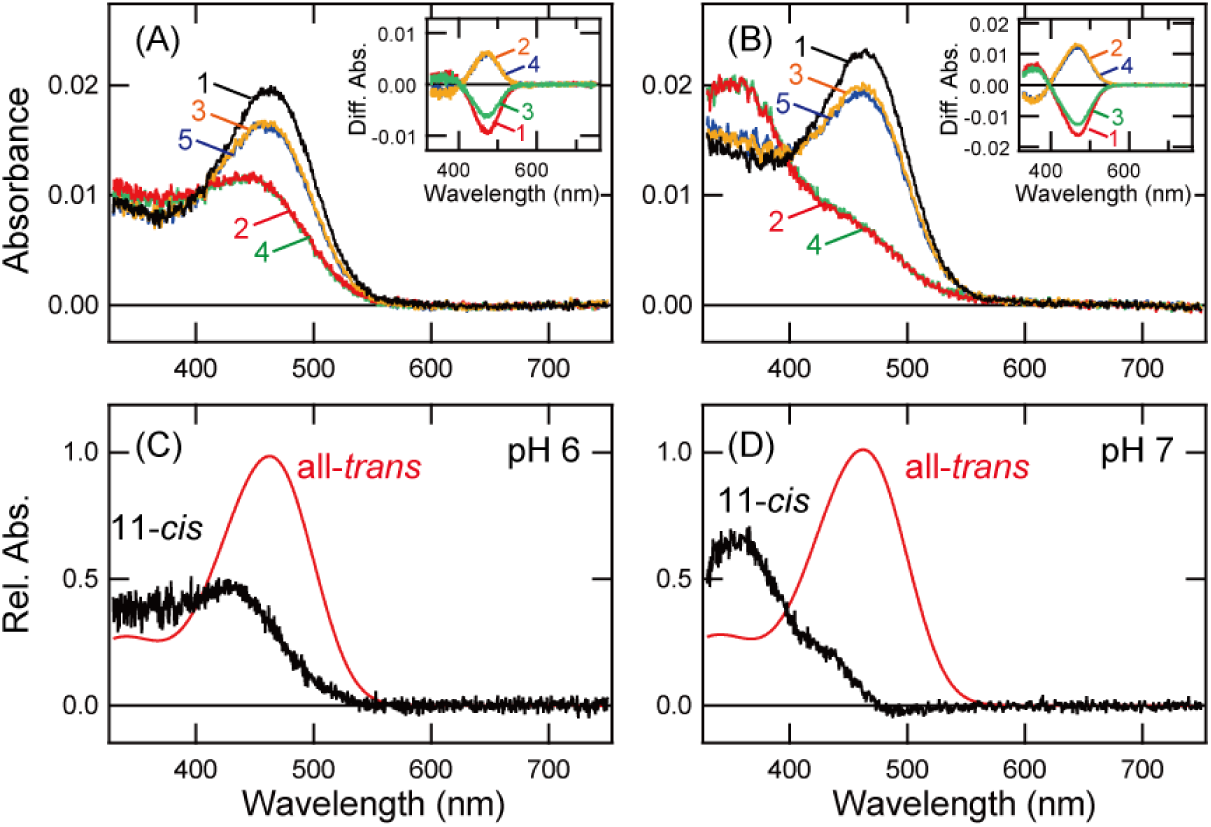
pH-dependent spectral changes of K91Y mutant of Opn5m. (A, B) Absorption spectra of K91Y mutant of Opn5m purified after the incubation with all-*trans* retinal. Spectra were recorded in the dark (curve 1), after yellow light irradiation (>500 nm) (curve 2), after subsequent UV light (360 nm) irradiation (curve 3), after yellow light re-irradiation (curve 4) and after UV light re-irradiation (curve 5) at pH 6 (A) or pH 7 (B). (inset) Spectral changes caused by yellow light irradiation (curve 1), subsequent UV light irradiation (curve 2), yellow light re-irradiation (curve 3) and UV light re-irradiation (curve 4). (C, D) Calculated absorption spectra of K91Y mutant of Opn5m at pH 6 (C) or pH 7 (D). The method of calculating the spectra of 11-*cis* (black curve) and all-*trans* retinal (red curve) bound forms is provided in Fig. S8.

## Discussion

Since the first characterization of Opn5m as a UV light sensor (8), we have been trying to identify the amino acid residue(s) that determines the UV light sensitivity of Opn5m. In this study, our mutational analysis in the extracellular side of Helix II revealed that Lys91 is an important residue for the acquisition of the UV light sensitivity. Our comprehensive mutational analysis at Lys91 unveiled that the introduction of a lysine, arginine or tyrosine residue at this position was crucial for the UV light sensitivity. This is consistent with the conservation of the lysine residue at this position of vertebrate Opn5m (Fig. S1A). The previous studies about Drosophila opsins showed that Lys90 is important for the UV light sensitivity (22, 23). Also, in UV-sensitive non-visual opsin, Opn3, found in a marine annelid, the lysine or arginine residue at position 94 contributes to the acquisition of the UV light sensitivity (24). Thus, these UV-sensitive opsins may have a similar molecular mechanism for the deprotonation of the Schiff base, that is, preventing the protonation of the Schiff base by the positive charge in the side chain of the residues in the extracellular side of Helix II. Moreover, introduction of a tyrosine residue at Lys91 produced a unique mixture of two components, one with a protonated and the other with a deprotonated Schiff base in Opn5m. Previous studies of vertebrate UV-sensitive cone pigments showed that the introduction of a serine, threonine, or tyrosine residue at positions 86 or 90 resulted in the formation of violet light-sensitive opsins (20, 25). In this study, K91T and K91S mutants of Opn5m were also violet light-sensitive opsins (Figs. 2, S4 and S5). Thus, an amino acid residue whose side chain contains a hydroxyl group would lead to the protonation of the Schiff base in these opsins. The different effects of the introduction of a tyrosine residue and serine/threonine residues in Opn5m may be due to different volumes of the side chains. The spatial proximity between the Schiff base and the residues in the extracellular side of Helix II and the hydrogen-bonding network connecting them would regulate the pKa of the Schiff base within the protein, which has been discussed in previous papers about vertebrate cone pigments (26, 27).

The mutations at Lys91 had drastic effects on not only the spectral property but also other molecular properties of Opn5m. The mutations did not hinder the bistable photoreaction but altered the preferential binding of the retinal isomers. Wild-type Opn5m directly binds both 11-*cis* and all-*trans* retinals (9, 28). This would be important for the formation of the photo-pigments not only in the retina, which has an 11-*cis* retinal supplying system, but also in other tissues such as brain, which lacks a sufficient supply of 11-*cis* retinal. Our mutational results showed that K91R mutant exclusively bound 11-*cis* retinal and K91T mutant exclusively bound all-*trans* retinal (Figs. 2 and 4). Previous reports showed that T94I mutant of bovine rhodopsin acquired the ability to directly bind all-*trans* retinal (29) and K94T mutant of UV-sensitive marine annelid Opn3 lost the ability to directly bind all-*trans* retinal (24). However, this phenotype of K94T mutant of marine annelid Opn3 contrasted with the decreased affinity for 11-*cis* retinal of K91T mutant of Opn5m. Thus, although it is a common concept that amino acid residues in the extracellular side of Helix II control the preferential binding of the retinal isomers, the detailed molecular mechanism of this control may differ among opsins. Our previous studies identified two amino acid residues (positions 168 and 188) that control the affinity for the retinal isomers (9, 28). It can be speculated that the amino acid residues located at distant positions around the retinal would finely control the affinity for the retinal isomers.

In conclusion, we identified an important spectral tuning site of UV-sensitive Opn5m. This residue also contributed to maintaining the affinities for both 11-*cis* and all-*trans* retinals. Vertebrate Opn5m, including human and mouse Opn5, has a common spectral property, UV light sensitivity, and provides one of the molecular mechanisms determining the short wavelength limit that a wide range of vertebrate species can detect (8, 9). Therefore, the conservation of Lys91 among vertebrate Opn5m proteins would be necessary to enable non-visual opsin Opn5m to work as the shortest wavelength sensor in various tissues, including the retina and brain.

## Acknowledgements

We thank Dr. Elizabeth Nakajima for critical reading of the manuscript. We also thank Prof. Robert S. Molday for the generous gift of a Rho1D4-producing hybridoma. This work was supported in part by Grants-in Aid for Scientific Research of MEXT to T.Y. (24K09530), Japan Science and Technology Agency (JST) CREST to T.Y. (JPMJCR1753), Japan Agency for Medical Research and Development (AMED) CREST to T.Y. (22gm1510007), a grant from the Takeda Science Foundation to T.Y., a grant from the Research Foundation for Opto-Science and Technology to T.Y. and a grant from The Naito Foundation to T.Y. C.F. is supported by a JSPS Research Fellowship for Young Scientists.

## Author Contributions

T.Y. designed the research. T.Y. K.A., K.F. and C.F. conducted the experiments. T.Y. K.A., K.F., C.F., H.O. and Y.S. analyzed the data. T.Y. wrote the manuscript with editing by all authors.

## Data availability

All data are available in the main text or the supplementary materials.

## Materials and Methods

### Preparation of Opn5 recombinant proteins

To improve the expression level of *X. tropicalis* Opn5m (GenBank accession number XM_002935990) recombinant protein, we truncated 21 amino acid residues from the C-terminus, as described in our previous reports (9, 28). The C-terminal truncated cDNA of *X. tropicalis* Opn5m was tagged with the epitope sequence of the anti-bovine rhodopsin monoclonal antibody Rho1D4 at the C-terminus and was introduced into the mammalian expression vector pCAGGS (30). Site-directed mutations were introduced using the seamless ligation cloning extract (SLiCE) method (31). The plasmid DNA was transfected into HEK293S cells using the calcium phosphate method. 5 μM 11-*cis* or all-*trans* retinal was added to the medium 24 h after transfection and the cells were kept in the dark until they were collected 48 h after transfection. The reconstituted pigments were extracted from cell membranes with 1 % DDM in Buffer A (50mM HEPES, 140mM NaCl, pH 6.5) and were purified using Rho1D4-conjugated agarose. The purified pigments were eluted with 0.02 % DDM in Buffer A containing the synthetic peptide that corresponds to the C-terminus of bovine rhodopsin. All of the procedures were carried out on ice under dim red light.

### Spectrophotometry

UV/Vis absorption spectra were recorded using a Shimadzu UV2400, UV2450 or UV2600 spectrophotometer and an optical cell (width, 2 mm; light path, 1 cm). The sample temperature was maintained using a temperature controller (RTE-210, NESLAB) at 0 ± 0.1 °C. The sample was irradiated with light which was generated by a 1-kW tungsten halogen lamp (Master HILUX-HR, Rikagaku seiki) and passed through optical filters (Y-52 or UV-D36C, AGC Techno Glass).

### HPLC analysis of retinal isomers

Retinal configurations within Opn5m samples were analyzed by HPLC (LC-10AT VP or LC-20AD; Shimadzu) equipped with a silica column (150×6 mm, A-012-2; YMC) as previously described (32).

## References

1. Shichida, Y., and Matsuyama, T. (2009) Evolution of opsins and phototransduction Philos Trans R Soc Lond B Biol Sci 364, 2881–2895

2. Ramirez, M. D., Pairett, A. N., Pankey, M. S., Serb, J. M., Speiser, D. I., Swafford, A. J. et al. (2016) The Last Common Ancestor of Most Bilaterian Animals Possessed at Least Nine Opsins Genome Biol Evol 8, 3640–3652

3. Hofmann, K. P., and Lamb, T. D. (2023) Rhodopsin, light-sensor of vision Prog Retin Eye Res 93, 101116

4. Yamashita, T. (2020) Unexpected molecular diversity of vertebrate nonvisual opsin Opn5 Biophys Rev 12, 333–338

5. Tarttelin, E. E., Bellingham, J., Hankins, M. W., Foster, R. G., and Lucas, R. J. (2003) Neuropsin (Opn5): a novel opsin identified in mammalian neural tissue FEBS Lett 554, 410–416

6. Sato, K., and Ohuchi, H. (2024) Molecular Property, Manipulation, and Potential Use of Opn5 and Its Homologs J Mol Biol 436, 168319

7. Tomonari, S., Migita, K., Takagi, A., Noji, S., and Ohuchi, H. (2008) Expression patterns of the opsin 5-related genes in the developing chicken retina Dev Dyn 237, 1910–1922

8. Yamashita, T., Ohuchi, H., Tomonari, S., Ikeda, K., Sakai, K., and Shichida, Y. (2010) Opn5 is a UV-sensitive bistable pigment that couples with Gi subtype of G protein Proc Natl Acad Sci U S A 107, 22084–22089

9. Yamashita, T., Ono, K., Ohuchi, H., Yumoto, A., Gotoh, H., Tomonari, S. et al. (2014) Evolution of mammalian Opn5 as a specialized UV-absorbing pigment by a single amino acid mutation J Biol Chem 289, 3991–4000

10. Buhr, E. D., and Van Gelder, R. N. (2014) Local photic entrainment of the retinal circadian oscillator in the absence of rods, cones, and melanopsin Proc Natl Acad Sci U S A 111, 8625–8630

11. Buhr, E. D., Vemaraju, S., Diaz, N., Lang, R. A., and Van Gelder, R. N. (2019) Neuropsin (OPN5) Mediates Local Light-Dependent Induction of Circadian Clock Genes and Circadian Photoentrainment in Exposed Murine Skin Curr Biol 29, 3478–3487 e3474

12. Buhr, E. D., Yue, W. W., Ren, X., Jiang, Z., Liao, H. W., Mei, X. et al. (2015) Neuropsin (OPN5)-mediated photoentrainment of local circadian oscillators in mammalian retina and cornea Proc Natl Acad Sci U S A 112, 13093–13098

13. Ota, W., Nakane, Y., Hattar, S., and Yoshimura, T. (2018) Impaired Circadian Photoentrainment in Opn5-Null Mice iScience 6, 299–305

14. Jiang, X., Pardue, M. T., Mori, K., Ikeda, S. I., Torii, H., D’Souza, S. et al. (2021) Violet light suppresses lens-induced myopia via neuropsin (OPN5) in mice Proc Natl Acad Sci U S A 118,

15. Nguyen, M. T., Vemaraju, S., Nayak, G., Odaka, Y., Buhr, E. D., Alonzo, N. et al. (2019) An opsin 5-dopamine pathway mediates light-dependent vascular development in the eye Nat Cell Biol 21, 420–429

16. Zhang, K. X., D’Souza, S., Upton, B. A., Kernodle, S., Vemaraju, S., Nayak, G. et al. (2020) Violet-light suppression of thermogenesis by opsin 5 hypothalamic neurons Nature 585, 420–425

17. Nakane, Y., Ikegami, K., Ono, H., Yamamoto, N., Yoshida, S., Hirunagi, K., et al. (2010) A mammalian neural tissue opsin (Opsin 5) is a deep brain photoreceptor in birds Proc Natl Acad Sci U S A 107, 15264-15268

18. Fukuda, A., Sato, K., Fujimori, C., Yamashita, T., Takeuchi, A., Ohuchi, H. et al. (2025) Direct photoreception by pituitary endocrine cells regulates hormone release and pigmentation Science 387, 43–48

19. Sato, K., Yamashita, T., Haruki, Y., Ohuchi, H., Kinoshita, M., and Shichida, Y. (2016) Two UV-Sensitive Photoreceptor Proteins, Opn5m and Opn5m2 in Ray-Finned Fish with Distinct Molecular Properties and Broad Distribution in the Retina and Brain PLoS One 11, e0155339

20. Fasick, J. I., Applebury, M. L., and Oprian, D. D. (2002) Spectral tuning in the mammalian short-wavelength sensitive cone pigments Biochemistry 41, 6860–6865

21. Koyanagi, M., Wada, S., Kawano-Yamashita, E., Hara, Y., Kuraku, S., Kosaka, S. et al. (2015) Diversification of non-visual photopigment parapinopsin in spectral sensitivity for diverse pineal functions BMC Biol 13, 73

22. Sakai, K., Tsutsui, K., Yamashita, T., Iwabe, N., Takahashi, K., Wada, A., et al. (2017) Drosophila melanogaster rhodopsin Rh7 is a UV-to-visible light sensor with an extraordinarily broad absorption spectrum Sci Rep 7, 7349

23. Salcedo, E., Zheng, L., Phistry, M., Bagg, E. E., and Britt, S. G. (2003) Molecular basis for ultraviolet vision in invertebrates J Neurosci 23, 10873–10878

24. Tsukamoto, H., Chen, I. S., Kubo, Y., and Furutani, Y. (2017) A ciliary opsin in the brain of a marine annelid zooplankton is ultraviolet-sensitive, and the sensitivity is tuned by a single amino acid residue J Biol Chem 292, 12971–12980

25. Yokoyama, S., Radlwimmer, F. B., and Blow, N. S. (2000) Ultraviolet pigments in birds evolved from violet pigments by a single amino acid change Proc Natl Acad Sci U S A 97, 7366–7371

26. Yokoyama, S., Starmer, W. T., Takahashi, Y., and Tada, T. (2006) Tertiary structure and spectral tuning of UV and violet pigments in vertebrates Gene 365, 95–103

27. Yokoyama, S., Tada, T., Liu, Y., Faggionato, D., and Altun, A. (2016) A simple method for studying the molecular mechanisms of ultraviolet and violet reception in vertebrates BMC Evol Biol 16, 64

28. Fujiyabu, C., Sato, K., Nishio, Y., Imamoto, Y., Ohuchi, H., Shichida, Y., et al. (2022) Amino acid residue at position 188 determines the UV-sensitive bistable property of vertebrate non-visual opsin Opn5 Commun Biol 5, 63

29. Ramon, E., del Valle, L. J., and Garriga, P. (2003) Unusual thermal and conformational properties of the rhodopsin congenital night blindness mutant Thr-94 --> Ile J Biol Chem 278, 6427–6432

30. Niwa, H., Yamamura, K., and Miyazaki, J. (1991) Efficient selection for high-expression transfectants with a novel eukaryotic vector Gene 108, 193–199

31. Motohashi, K. (2017) Seamless Ligation Cloning Extract (SLiCE) Method Using Cell Lysates from Laboratory Escherichia coli Strains and its Application to SLiP Site-Directed Mutagenesis Methods Mol Biol 1498, 349–357

32. Tsutsui, K., Imai, H., and Shichida, Y. (2007) Photoisomerization efficiency in UV-absorbing visual pigments: protein-directed isomerization of an unprotonated retinal Schiff base Biochemistry 46, 6437–6445

