## Supplementary information for "A key spectral tuning site of UV-sensitive vertebrate non-visual opsin Opn5"

(A)

| | 86 90 91 |
|---|---|
| Bovine Rhodopsin | AVADLFMVFGGFTTTLYTSL |
| Homo sapiens Opn5 | AVCDLGISVVGKPFTIISCF |
| Mus musculus Opn5 | AVCDLGISVVGKPFTIISCF |
| Gallus gallus Opn5m | AVCDLGISVVGKPFSIISFF |
| Anolis carolinensis Opn5m | AVCDLGISVVGKPFSIIAFF |
| Chrysemys picta Opn5m | AVCDLGISVVGKPFSIISFF |
| Xenopus tropicalis Opn5m | AVCDLGISVTGKPFAIVSCF |
| Ambystoma mexicanum Opn5m | AFCDLGISVVGKPFIIASFF |
| Lepisosteus oculatus Opn5m | AIFDFGISVTGKPFFVISSF |
| Danio rerio Opn5m | AIFDFGISVSGKPFFIVSSF |
| Oryzias latipes Opn5m | AIFDFGISVTGKPFFVVSSF |

(B)

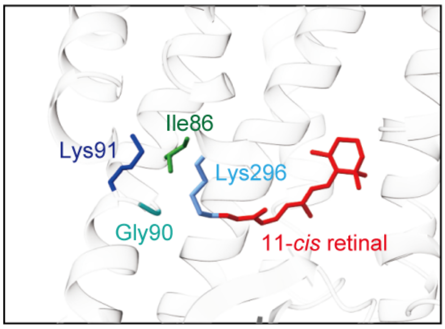


**Figure S1 Amino acid residues at positions 86, 90 and 91 in vertebrate Opn5m**

(A) Comparison of the amino acid residues in the extracellular side of Helix II among bovine rhodopsin and various vertebrate Opn5m proteins. Amino acid sequences of bovine rhodopsin (GenBank accession number: K00506), human Opn5 (AY377391), mouse Opn5 (AY318865), chicken Opn5m (AB368182), anole lizard Opn5m (XM_003215359), painted turtle Opn5m (XM_005297358), *Xenopus tropicalis* Opn5m (XM_002935990), Mexican salamander Opn5m (XM_069605841), spotted gar Opn5m (XM_015349763), zebrafish Opn5m (AY493740) and medaka fish Opn5m (XM_023953178) are shown. The three residues at positions 86, 90 and 91 are highlighted in red. Amino acids are numbered based on the bovine rhodopsin numbering system. (B) Predicted structural arrangement of the residues at positions 86, 90 and 91 of human Opn5. Structural model of human Opn5 was constructed based on the crystal structure of bovine rhodopsin (PDB: 1U19) by homology modeling using MOE software (Chemical Computing Group Inc.) and was visualized using UCSF ChimeraX.

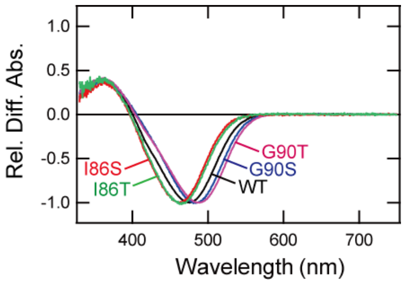


**Figure S2 Comparison of spectral property of I86 and G90 mutants**

Difference spectra shown as curve 1 in Fig. 1C-1G were normalized to be ~ -1.0 at the negative maximum.

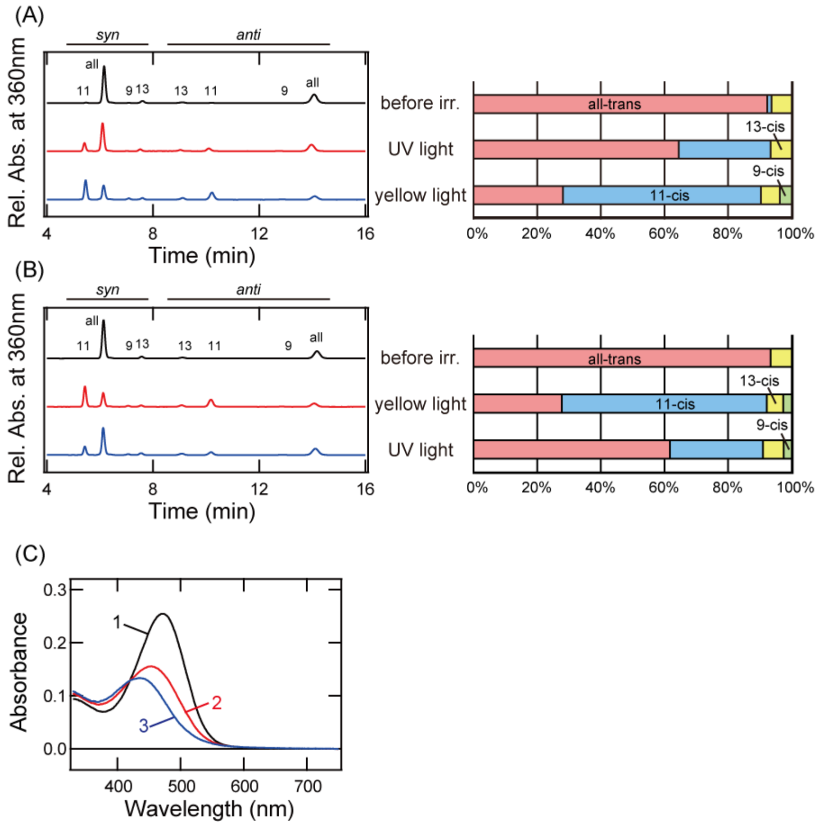


**Figure S3 Characterization of K91T mutant of Opn5m**

(A) Retinal configuration changes of K91T mutant purified after the incubation with 11-*cis* retinal. The configurations were analyzed before light irradiation, after UV light (360 nm) irradiation and after subsequent yellow light (>500 nm) irradiation. (B) Retinal configuration changes of K91T mutant purified after the incubation with all-*trans* retinal. The configurations were analyzed before light irradiation, after yellow light irradiation and after subsequent UV light irradiation. (Left) The retinal configurations were analyzed with HPLC after extraction of the chromophore as retinal oximes (syn and anti forms of 9-*cis*, 11-*cis*, 13-*cis*, and all-*trans* retinal oximes). (Right) Isomeric compositions of retinal before and after light irradiation of the mutant. (C) Calculation of absorption spectra of K91T mutant. We recorded the absorption spectrum (curve 1) of the sample purified after reconstitution with all-*trans* retinal (the same as curve 1 of Fig. 2B). This sample contained almost exclusively all-*trans* retinal (see Fig. S3B), which means that curve 1 corresponds to the spectrum of all-*trans* retinal bound form. To obtain the absorption spectrum of the 11-*cis* retinal bound form (curve 3), we subtracted curve 1 from curve 2 (the same as curve 2 of Fig. 2B) based on the component ratio of 11-*cis* and all-*trans* retinals shown in Fig. S3B. Finally, we normalized curves 1 and 3 to be ~1.0 at λmax of curve 1 to show the normalized spectra in Fig. 2C.

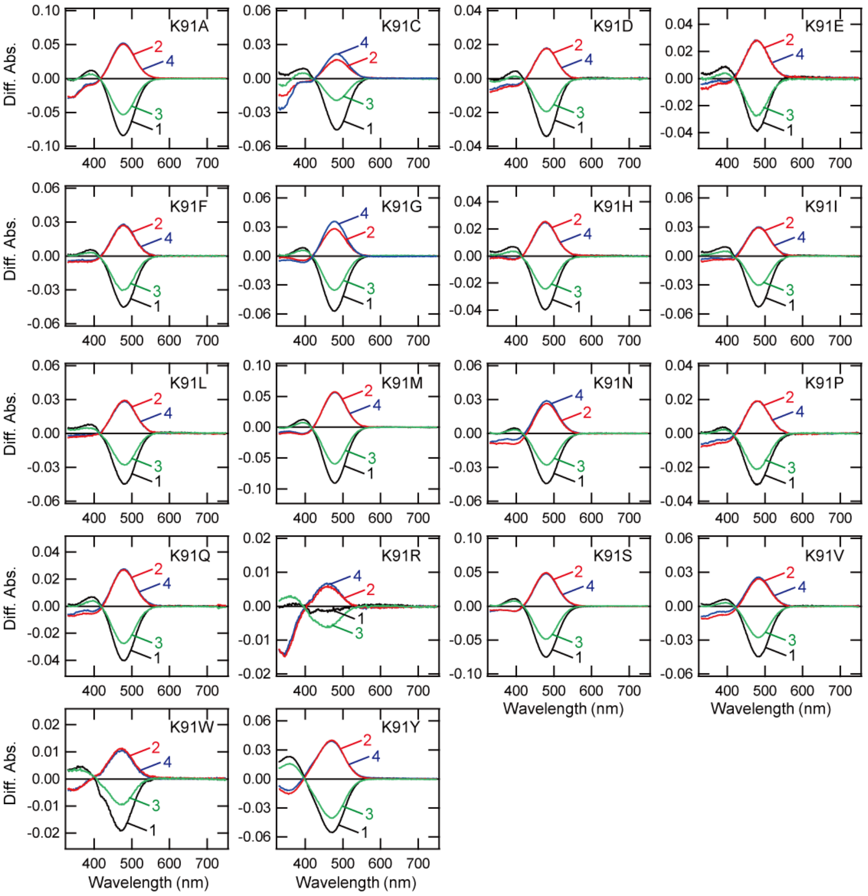


**Figure S4 Spectral property of K91 mutants after the incubation with all-*trans* retinal**

The cell membranes containing K91 mutants after the addition of all-*trans* retinal were solubilized with 1 % DDM and their absorption spectra were recorded in the dark and after light irradiation. Spectral changes caused by yellow light (>500 nm) irradiation (curve 1), subsequent UV light (360 nm) irradiation (curve 2), yellow light re-irradiation (curve 3) and UV light re-irradiation (curve 4) are shown.

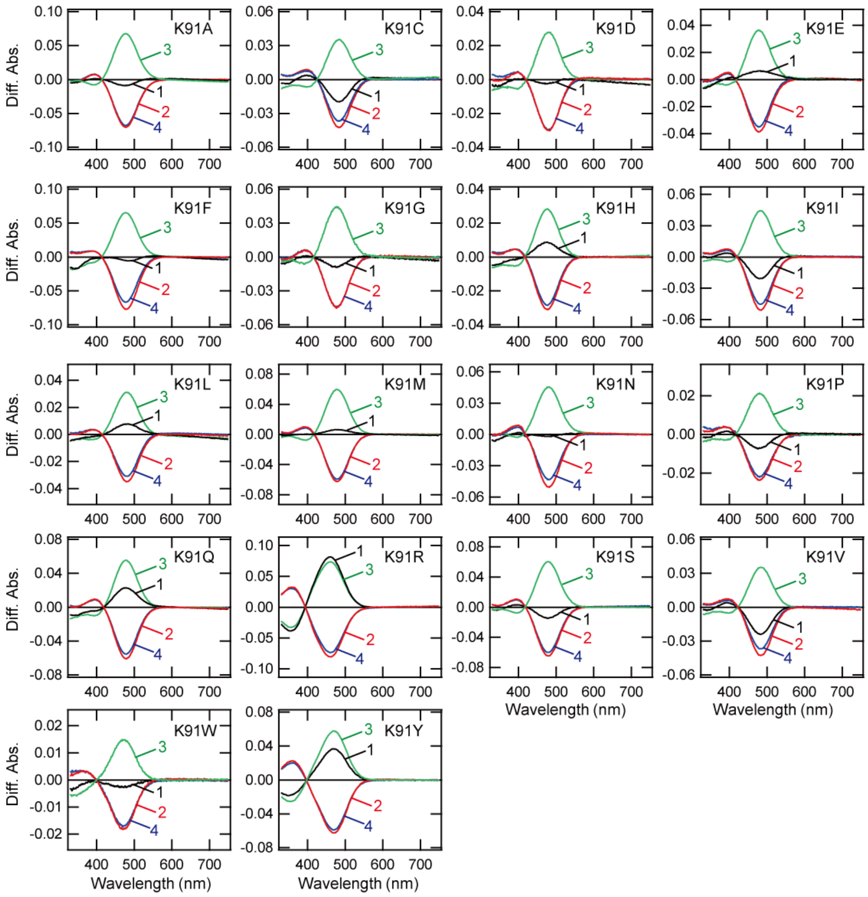


**Figure S5 Spectral property of K91 mutants after the incubation with 11-*cis* retinal**

The cell membranes containing K91 mutants after the addition of 11-*cis* retinal were solubilized with 1 % DDM and their absorption spectra were recorded in the dark and after light irradiation. Spectral changes caused by UV light (360 nm) irradiation (curve 1), subsequent yellow light (>500 nm) irradiation (curve 2), UV light re-irradiation (curve 3) and yellow light re-irradiation (curve 4) are shown.

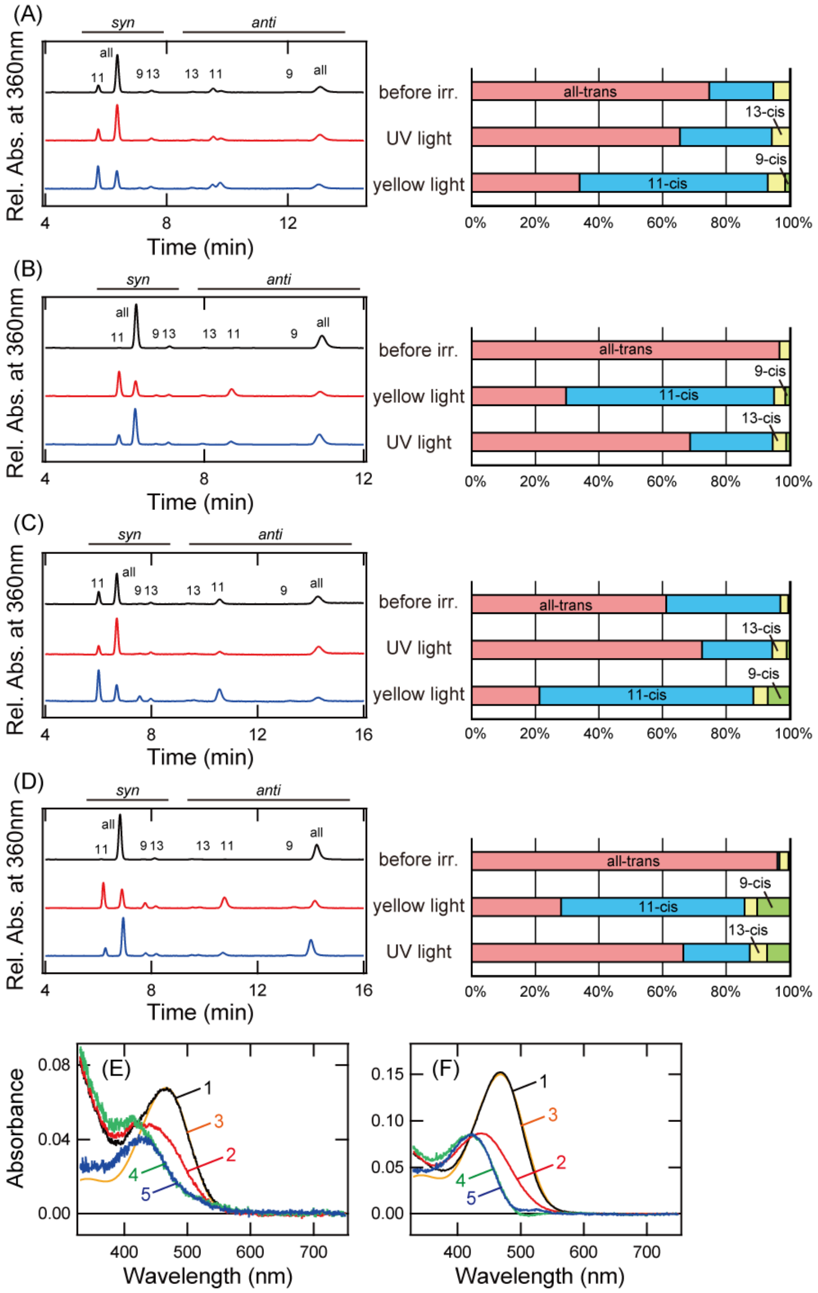


**Figure S6 Characterization of K91A and K91Q mutants of Opn5m**

(A, C) Retinal configuration changes of K91A (A) and K91Q (C) mutants purified after the incubation with 11-*cis* retinal. The configurations were analyzed before light irradiation, after UV light (360 nm) irradiation and after subsequent yellow light (>500 nm) irradiation. (B, D) Retinal configuration changes of K91A (B) and K91Q (D) mutants purified after the incubation with all-*trans* retinal. The configurations were analyzed before light irradiation, after yellow light irradiation and after subsequent UV light irradiation. (Left) The retinal configurations were analyzed with HPLC after extraction of the chromophore as retinal oximes (syn and anti forms of 9-*cis*, 11-*cis*, 13-*cis*, and all-*trans* retinal oximes). (Right) Isomeric compositions of retinal before and after light irradiation of the mutant. (E, F) Calculation of absorption spectra of K91A (E) and K91Q (F) mutants. We recorded the absorption spectrum (curve 1) of the samples purified after reconstitution with all-*trans* retinal (the same as curve 1 of Fig. 3B or 3D). This sample almost exclusively contained all-*trans* retinal (see Fig. S6B or S6D). To obtain the absorption spectrum of the 11-*cis* retinal bound form (curve 4), we subtracted curve 1 from curve 2 (the same as curve 2 of Fig. 3B or 3D) based on the component ratio of 11-*cis* and all-*trans* retinals shown in Fig. S6B or S6D. To remove the light scattering component in curves 1 and 4, we fitted curve 1 with a template spectrum modeled by the Lamb and Govardovskii method (1, 2) (curve 3). We also subtracted the difference spectrum between curves 1 and 3 from curve 4 (curve 5). Finally, we normalized curves 3 and 5 to be ~1.0 at λmax of curve 3 to show the normalized spectra in Fig. 3E or 3F.

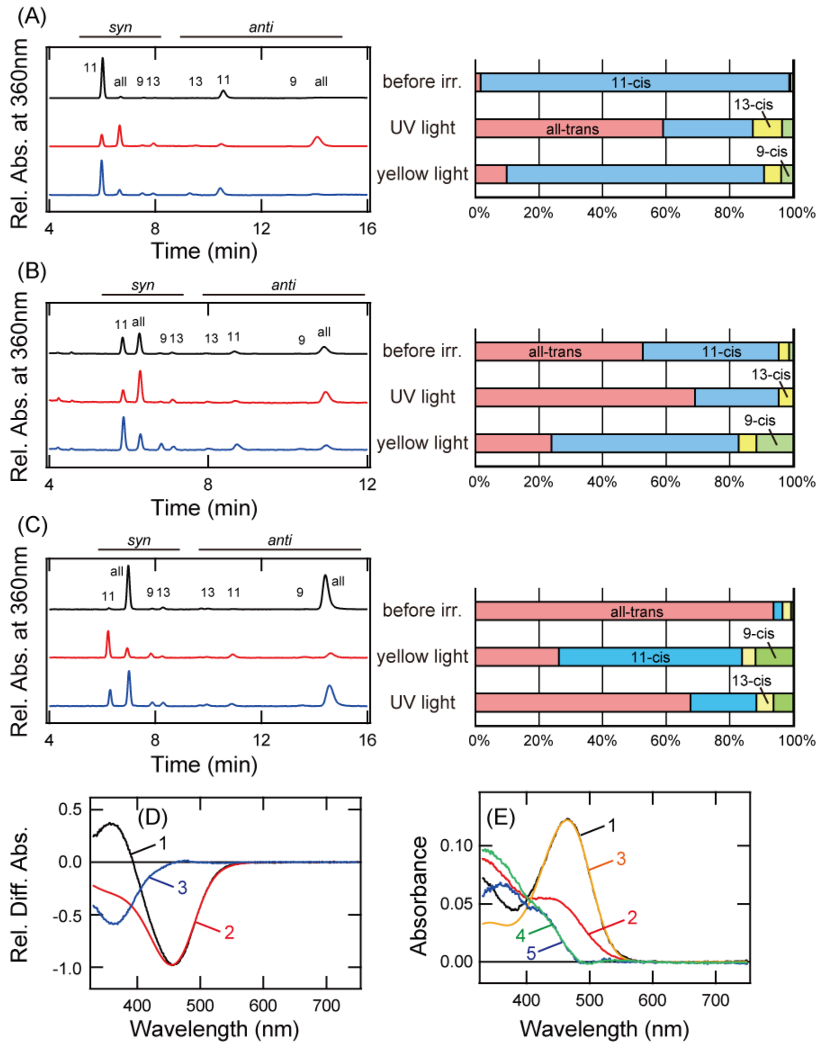


**Figure S7 Characterization of K91R and K91Y mutants of Opn5m**

(A, B) Retinal configuration changes of K91R (A) and K91Y (B) mutants purified after the incubation with 11-*cis* retinal. The configurations were analyzed before light irradiation, after UV light (360 nm) irradiation and after subsequent yellow light (>500 nm) irradiation. (C) Retinal configuration changes of K91Y mutant purified after the incubation with all-*trans* retinal. The configurations were analyzed before light irradiation, after yellow light irradiation and after subsequent UV light irradiation. (Left) The retinal configurations were analyzed with HPLC after extraction of the chromophore as retinal oximes (syn and anti forms of 9-*cis*, 11-*cis*, 13-*cis*, and all-*trans* retinal oximes). (Right) Isomeric compositions of retinal before and after light irradiation of the mutant.

(D) Calculation of absorption spectra of K91R mutant. We normalized the difference spectrum (curve 2 in the inset of Fig. 4A) to be ~ -1.0 at the negative maximum (curve 1). To obtain the spectrum of the all-*trans* retinal bound form (curve 2), we fitted curve 1 with a template spectrum modeled by the Lamb and Govardovskii method (1, 2). To obtain the spectrum of the 11-*cis* retinal bound form (curve 3), we subtracted curve 1 from curve 2. Finally, we normalized curves 2 and 3 to be ~1.0 at the negative maximum of curve 2 to show the normalized spectra in Fig. 4E. (E) Calculation of absorption spectra of K91Y mutant. We recorded the absorption spectrum (curve 1) of the sample purified after reconstitution with all-*trans* retinal (the same as curve 1 of Fig. 4D). This sample almost exclusively contained all-*trans* retinal (see Fig. S7C). To obtain the absorption spectrum of the 11-*cis* retinal bound form (curve 4), we subtracted curve 1 from curve 2 (the same as curve 2 of Fig. 4D) based on the component ratio of 11-*cis* and all-*trans* retinals shown in Fig. S7C. To remove the light scattering component in curves 1 and 4, we fitted curve 1 with a template spectrum modeled by the Lamb and Govardovskii method (1, 2) (curve 3). We also subtracted the difference spectrum between curves 1 and 3 from curve 4 (curve 5). Finally, we normalized curves 3 and 5 to be ~1.0 at $\lambda max$ of curve 3 to show the normalized spectra in Fig. 4F.

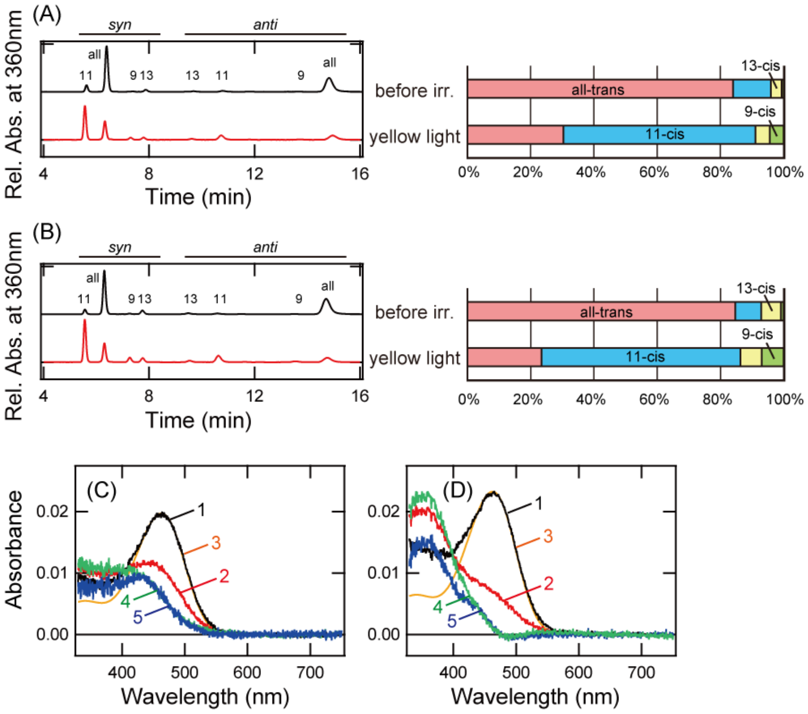


**Figure S8 Characterization of K91Y mutants under pH 6 and 7**

(A, B) Retinal configuration changes of K91Y mutant purified after the incubation with all-*trans* retinal. The configurations were analyzed before light irradiation and after subsequent yellow light (>500 nm) irradiation under pH 6 (A) and pH 7 (B). (Left) The retinal configurations were analyzed with HPLC after extraction of the chromophore as retinal oximes (syn and anti forms of 9-*cis*, 11-*cis*, 13-*cis*, and all-*trans* retinal oximes). (Right) Isomeric compositions of retinal before and after light irradiation of the mutant. (C, D) Calculation of absorption spectra of K91Y mutant under pH 6 (C) and pH 7 (D). We recorded the absorption spectrum (curve 1) of the samples purified after reconstitution with all-*trans* retinal (the same as curve 1 of Fig. 5A or 5B). This sample almost exclusively contained all-*trans* retinal (see Fig. S8A or S8B). To obtain the absorption spectrum of the 11-*cis* retinal bound form (curve 4), we subtracted curve 1 from curve 2 (the same as curve 2 of Fig. 5A or 5B) based on the component ratio of 11-*cis* and all-*trans* retinals shown in Fig. S8A or S8B. To remove the light scattering component in curves 1 and 4, we fitted curve 1 with a template spectrum modeled by the Lamb and Govardovskii method (1, 2) (curve 3). We also subtracted the difference spectrum between curves 1 and 3 from curve 4 (curve 5). Finally, we normalized curves 3 and 5 to be ~1.0 at λmax of curve 3 to show the normalized spectra in Fig. 5C or 5D.

### References


1. Lamb, T. D. (1995) Photoreceptor spectral sensitivities: common shape in the long-wavelength region *Vision Res* **35**, 3083-3091
2. Govardovskii, V. I., Fyhrquist, N., Reuter, T., Kuzmin, D. G., and Donner, K. (2000) In search of the visual pigment template *Vis Neurosci* **17**, 509-528
